# Social and spatial drivers of the multitiered structure of zebra finch social networks

**DOI:** 10.1101/2024.11.25.625155

**Authors:** Yixuan Zhang, Xinyi Jiang, Lucy M. Aplin, Daiping Wang, Damien R. Farine

## Abstract

Social network structure plays a key role in shaping processes in animal populations. These networks often show distinct patterns in humans and other large mammals, with relationship strengths organized into different tiers. Here, we used continuous, fine-scale tracking of four large captive colonies of zebra finches (*Taeniopygia guttata*), revealing that zebra finches consistently have 1-2 closest contacts, 6-7 close contacts, and 22-24 strong contacts. The identities of these contacts remain stable across days, with strong contacts maintained by spatial affinity while closest and close contacts are maintained by social choice. These results suggest that zebra finches egocentric networks and social structure are made up of consistent, differentiated relationships forming a multitiered social structure.

The similarities in patterns to other species suggest that fundamental principles, such as limitations in time and the ability to move through social space, could drive common structural properties in animal social networks.

## Main

## Introduction

Social network structures have important implications for animal populations. They can shape the spread of disease (*1*), what novel behaviours emerge (*2*, *3*), the patterning of cultures (*4*), and whether individuals are likely to cooperate or not (*5*, *6*). Thus, how localised interactions scale up to shape the patterns of connections among individuals in a population is a central question in social evolution (*7*). The importance of social relationships (*8*, *9*) is reflected in the diversity of ways that social structure can be expressed. First, individuals can have multiple different types of interactions with a given social partner (e.g. grooming, aggressive, play) (*10*). How these combine to form relationships is typically captured using multilayered or multiplex networks, whereby each layer represents one interaction type (*11*, *12*). Second, individuals can have different types of social partners (e.g. mate, alliance member, casual associate, competitor) (*13*), resulting in a variety of social relationships (*14*, *15*). Research on human societies has revealed distinct and consistent patterns in terms of the number and identity of individuals we interact with socially (*16*). From the frequency of calls (*17*) to co-authorship networks (*18*), individuals maintain a substantially higher rate of contact with a few closest social partners (*16*). More formally, human egocentric networks have a fractal structure that is characterised by an increasing number of social associates of decreasing relationship strength (with a consistent scaling ratio). Typically, relationship strengths can be classed into tiers representing 1-2 closest social partners, 5 close relationships, 15 strong relationships, and so on (*16*) [but see (*19*) for a review of the limitations of these findings]. Recent investigations into animal societies suggest that many species, including primates (*20–22*), cetaceans (*23*, *24*), other large terrestrial mammals (*25*, *26*), and birds (*27*, *28*), also live in societies defined by such differentiated relationships.

When aggregated across multiple individuals, differentiated social relationships can give rise to a multitiered—or multilevel—network structure (*16*, *29*). These are defined as having communities (sets of strongly connected individuals) that are composed of subcommunities (subsets of more strongly connected individuals, i.e. close and closest relationships). Such with the same close social partners, strong social partners (and so on). In some societies, individuals can freely join or leave social groups, even if they are commonly observed with the same set of associates (their closest social partners). An example of this is bottlenose dolphin (*Tursiops aduncus*) societies (*30*). In an extreme case, known as multilevel societies (*31*), subcommunities move and make social decisions as one cohesive social entity or social unit. Examples of stable social units are the one-male multi-female units that interact within some primate societies (*32*) or stable groups of vulturine guineafowl (*Acryllium vulturinum*) that fission-fusion with other stable groups (*28*). In both cases—those in which individuals make social choices and those where groups make social choices—can generate a multitiered social structure. However, while evidence for true multilevel societies is increasing rapidly (*31*), evidence for multitiered social structure arising in species where individuals make more autonomous decisions remains scarcer.

Detecting multitiered social structures in non-human animals is challenging. This is because it requires precisely knowing social preferences at the individual-level and tracking these over time to establish how stable they are. It then requires being able to obtain this information across a sufficient number (and complete set) of interconnected individuals to obtain a meaningful picture of population structure (*33*). Traditional sampling methods, such as visual observations of interactions among individuals, do not easily scale to large numbers of individuals (*34*) and thus have limited the ability of studies to capture the large-scale structural consequences of individual-level social decisions (Table 1). Such observations are especially challenging to conduct in birds. For this reason, the social lives of birds beyond cooperative breeding have been understudied, resulting in a mammal bias despite birds being potentially rich in complex social behaviours (*27*, *35*).

**Table 1.**
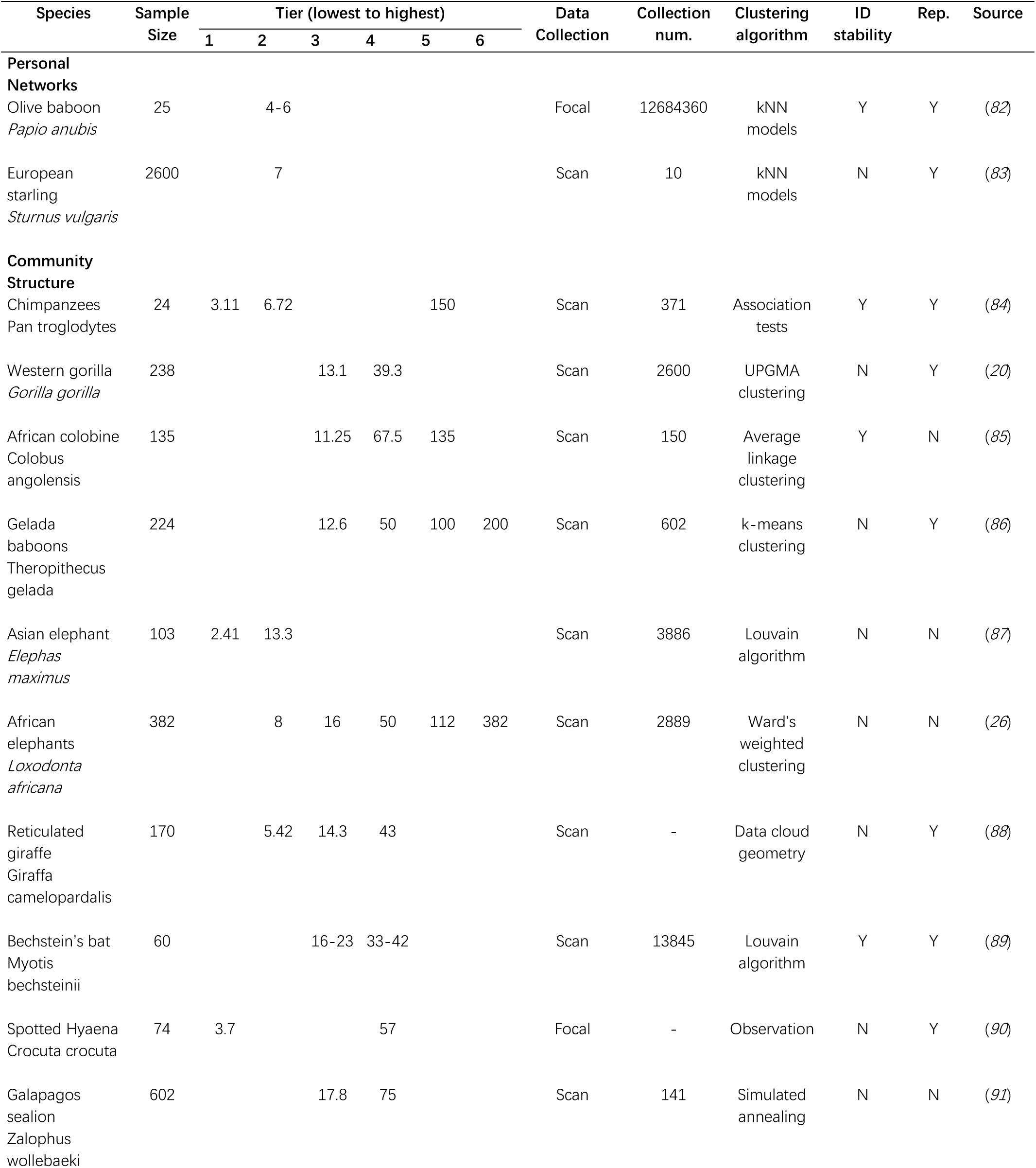
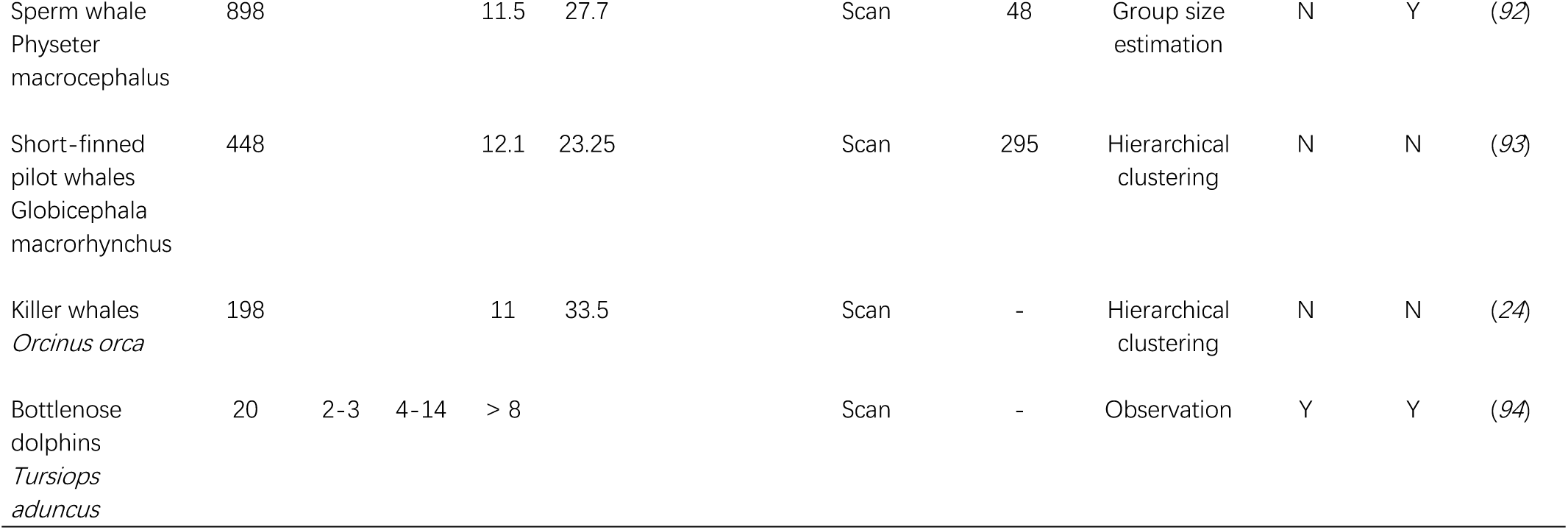
Mean tier sizes for non-human animals. (tier 1: small alliance or mother-offspring unit; tier 2: family; tier 3: bond group; tier 4: clan; tier 5: sub-population; tier 6: population). There is a scarcity of research on egocentric networks that detect the fractal hierarchical pattern of relationship differentiation. Similarly, studies investigating multitiered (including multilevel) social structures with extensive data collection, stable individual identities in groups, and experiment replication are also limited.

There are good reasons to believe that fractal patterns of relationships in egocentric networks, and emergent multitiered structures in larger-scale networks, may be more widespread than just in humans and large mammals (*27*, *36*). We predict that these are especially likely to be discovered in species that maintain long-term social bonds with both mates and non-reproductive social partners within a large, seemingly open society. This is because social relationships in animals take time to form (*37*). After all, individuals are generally limited in how many social interactions they can have at once (e.g. grooming relationships are generally dyadic) (*38*), and because the inherent embedding of social associations in space (*39*) promotes repeated associations within consistent subsets of individuals. Such drivers may, therefore, control the number of tiers (number of different types of social relationships) and community sizes of different tiers. A key question is whether they do so in similar ways across different species, and to what extent they are driven by individual social choice (*14*, *15*).

Here, we bridge the gap between studies of individual-level decision-making and population- level multitiered social structure by using continuous and fine-scale tracking in four large (N≍80, see table S1), replicated colonies of captive zebra finches (*Taeniopygia guttata*).

Zebra finches are highly social birds that can naturally form colonies of 24 to over 200 individuals (*40*, *41*). Within colonies, individuals typically move in mixed-sex pairs or small groups (*41*) and socialise at ‘social hotspots’ (*42*). Individuals form and maintain long-lasting non-breeding social preferences with other individuals (*43*), and these are often expressed by frequent ‘clumping’—two or more birds sitting on a perch in body contact with each other (*44*) where they allopreen or express other affiliative behaviours (*45*). Previous studies suggest that these colonies form the substrate for song cultures to emerge and be maintained (*46*) and maintain consistent group-level social structures (*47*); however, the mechanisms underlying how social structure emerges are still poorly understood (*48–50*).

In our study, each individual in each colony was fitted with a unique machine-readable barcode (*51*), allowing their position to be tracked continuously (every two seconds from morning to evening) over 36 days per colony. Birds were tracked after being introduced to unfamiliar members of the opposite sex and studied in mixed-sex colonies (e.g. N=40 males and 40 females). We then used the tracking data to construct egocentric and colony-level social contact networks (fig. S1, movie S1). A major challenge to drawing firm conclusions about how social structure emerges is that studies (i) typically confound spatial and social processes and (ii) use a wide range of different methods (Table 1). Here, we used a network edge definition that disentangles social choice from spatial co-occurrences (*52*) and null models (*53*) to determine the relative contributions of social and spatial choices on the structure of zebra finch social networks. Specifically, we quantified social contacts (i.e. ’clumping’) based on the proximity of barcodes (equivalent to two halves of a zebra finch body width apart) and defined network edge weights as the number of frames that two individuals were clumping divided by the number of frames in which both birds were located on the same perch (i.e. in the same spatial location).

Our approach allowed us to examine the decisions that individuals made—in terms of whom to associate with—conditioned on spatial proximity (i.e. being on the same perch) instead of confounding preferences for whom to associate with where in space individuals choose to be. Clumping behaviour never happens (or at least is not maintained) without social choice, as zebra finches maintain consistent inter-individual spacing on perches (see fig. S1 and movie S1). We then constructed two null models, one that randomised the identities of birds within perch and a second that randomised the identities of birds within the whole aviary.

These models allowed us to examine how observed results deviate from the expectations when removing only the preference for whom to clump with (within perch randomisations) or removing both social and spatial preferences (within aviary randomisations). To analyse the social networks, we used established methods from the human literature for egocentric networks (*54*) and methods recently proposed to test for the presence of multilevel social structures in animals (*31*). These methods are largely parameter-free, allowing us to objectively generate results that are directly comparable to current and future studies.

## Results

### Zebra finches have consistent closest, close, and strong social partners

The large amount of positional data (mean=1.88 million detections per aviary per day) enabled us to construct daily social networks. This allowed for both within- and between- colony replications when examining the consistency of egocentric network structure across days. We first used the Jenks natural breaks algorithm (*55*) to identify different categories of social contacts in the egocentric networks of 152 male and 151 female zebra finches (e.g. Fig. 1A, B). This algorithm identifies clusters by minimising within-cluster distances and maximising between-cluster distances. As expected, we found an increase in the goodness of variance fit (GVF) index with the number of clusters in each individual’s networks (i.e. GVF=1 when all the nodes are in the same cluster). We then used the cut-off at GVF = 0.85 ego-networks (N=303), the GVF indexes exceeded the recommended cut-off when the number of clusters was four (Fig. 1C). These results suggest that birds had four clusters (or tiers) of relationships: closest, close, and strong social partners, with the fourth cluster representing loose acquaintances or individuals without a strong social relationship (Table 2, fig. S3-6). The number of partners of each type did not differ between males and females (fig. S2A) and was consistent over time (Fig. 1D). Birds generally had more opposite-sex social partners of all types. However, females had more same-sex closest and close partners compared to males, while males had more opposite-sex partners compared to females (fig. S2B).

**Figure 1.**
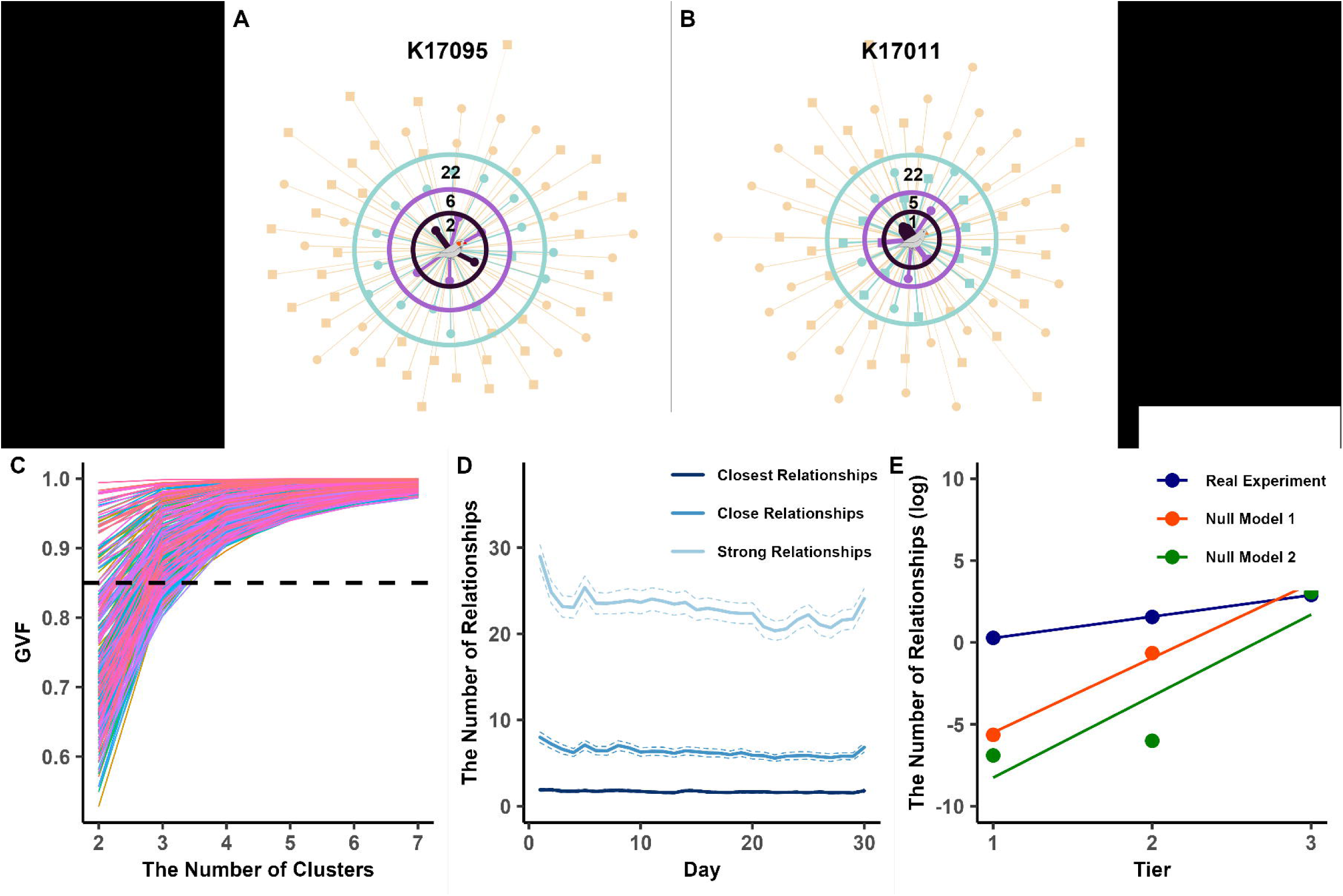
The fractal social structure in zebra finches using Jenks natural breaks algorithm in from four large(n=80) replicated colonies. Observed egocentric network in a (A) male and (B) female, showing closest (inner circle), close (middle circle), and strong (outermost circle) relationships (squares=males, circles=females) based on Jenks natural breaks algorithm in real experiments’ raw data. (C) Goodness of variance fit (GVF) index for Jenks natural breaks algorithm for daily ego-network of 303 individuals, with GVF = 0.85 (dashed line) consistently revealing four clusters in real experiments. (D) The number and temporal trend of social partners in each cluster was largely consistent across individuals and across time. The solid lines represent mean value, and the dashed lines show 95% confidence intervals. (E) The number of preferred social partners increased exponentially across different tiers. The points are the logarithmic mean value of the number of relationships.

**Table 2.** The number of relationships across different tiers among the real experiment and the two null models. Values in brackets represent 95% confidence intervals (n = 303).

| Tier | Number of Relationship |  |  |
| --- | --- | --- | --- |
|  | Real Experiment | Null Model 1 | Null Model 2 |
| 1 (lowest) | 1.69 (1.63-1.75) | 0.25 (0.21-0.28) | 0.00 (0.00-0.01) |
| 2 | 6.30 (6.06-6.54) | 4.09 (3.85-4.33) | 1.00 (0.79-1.22) |
| 3 | 22.97 (22.26-23.68) | 40.85 (39.94-41.76) | 50.60 (49.43-51.78) |
| 4 (highest) | 51.63 (50.55-52.72) | 33.75 (32.69-34.81) | 24.00 (22.78-25.22) |

We then tested for the occurrence of a fractal pattern of relationship differentiation in the egocentric networks. Fractal patterns occur when the number of preferred social partners increases exponentially across different tiers, resulting in a consistent ratio in the number of social partners from one tier to the next (*16*). This analysis revealed that the number of relationships at higher tiers is 3.77 times (e^1.326, the slope of the regression equation) that of the tier below it (Fig. 1E). This linear slope of the log of the number of relationships in each tier provides evidence that zebra finch egocentric social networks are structured in a manner that is consistent with fractal patterns.

The strength of an individual’s contact with each member of the colony was also consistent from one day to the next. The average cosine similarity (*57*) across individuals ranged from 0.67 to 0.76 (full results in table S2), which is substantially higher than what is expected by chance (0.19-0.35, calculated by randomly swapping the identities of individuals within each network but differently across days). This suggests that not only is the structure of the egocentric networks consistent from day to day but also the identities of the closest, close and strong social partners are consistent from one day to the next.

To gain a better understanding of the relative roles of spatial co-occurrence and individual social choices in determining the distribution of social relationships in egocentric networks, we employed two null models. In the first null model, we tested for major social preferences by permuting the identities of individuals on the same perch at the same time. This examines the consequences of individual decisions about who to have body contact with within the subset of most likely (or spatially proximate) associates. We note that the presence of two individuals on the same perch may also represent a social preference. However this would unlikely to result in clumping by chance. In the second null model, we tested for the effect of spatial preferences (which perch to use) on who individuals interact with by permuting the individual identities across perches.

We found that when removing spatial and social preferences (null model 2), individuals maintained no closest and close relationship (Table 2). Of note, the relationship between tier and the log of number of social partners in each tier was not linear, indicating a loss of the fractal nature of the egocentric networks under null model 2 (Fig. 1E). When maintaining spatial, but no social, choices (null model 1), individuals still expressed closest and close relationships, but fewer than in the observed data (Table 2). Under null model 1, the slope for the regression suggests a reasonable fit to the data (Fig. 1E). However the slope is very steep, indicating that the number of relationships grows disproportionately faster at higher tiers (e.g. from 0.2 to 4, and from 4 to 40). Finally, it was only in the observed data that we detected all three lower (or inner) tiers (containing < 40 individuals, Table 2). Together, these models suggest that similarities in space use play an important role in partitioning societies into broad social units (i.e. forming the higher, or outer, tiers of preferences in egocentric networks, Fig. 1E), which are commonly known as communities, but that the social decisions that individuals make within their spatial associates play a critical role in generating lower tiers, in which individuals express more distinct social preferences (i.e. more relationships with higher strength were shown in real data, Fig. 1E).

### Consistent, differentiated relationships result in multitiered social networks

Individual zebra finches express clear social preferences within their most spatially proximate conspecifics. These patterns should not—in and of themselves—produce a multitiered structure. For example, territorial individuals have a distinct set of first-degree neighbours, second-degree neighbours, and so on. However, the overall structure is lattice- like, meaning that second-degree neighbours are rarely shared among first-degree neighbours. This contrasts with clustered sets of strongly interconnected individuals that share similar closest, close, and strong social partners, forming a multitiered social structure.

To test whether the social preferences of zebra finches resulted in a multitiered social structure, we applied the ’Louvain’ community detection algorithm, which uses an iterative approach to identify hierarchical subgroups by maximising connection density within versus between groups across tiers (*31*). This analysis revealed second and third tier communities in each colony (the first tier represents each single individual as a distinct community), with bootstrapping analyses (*56*) confirming that this structure was robust (Table 3). The average size of the second tier community was 3.68 and 8.55 for the third tier community (Fig. 2A, table S3). These values are slightly larger than the closest and close associates found using the egocentric network approach (by one individual in each, after accounting for the ego).

**Figure 2.**
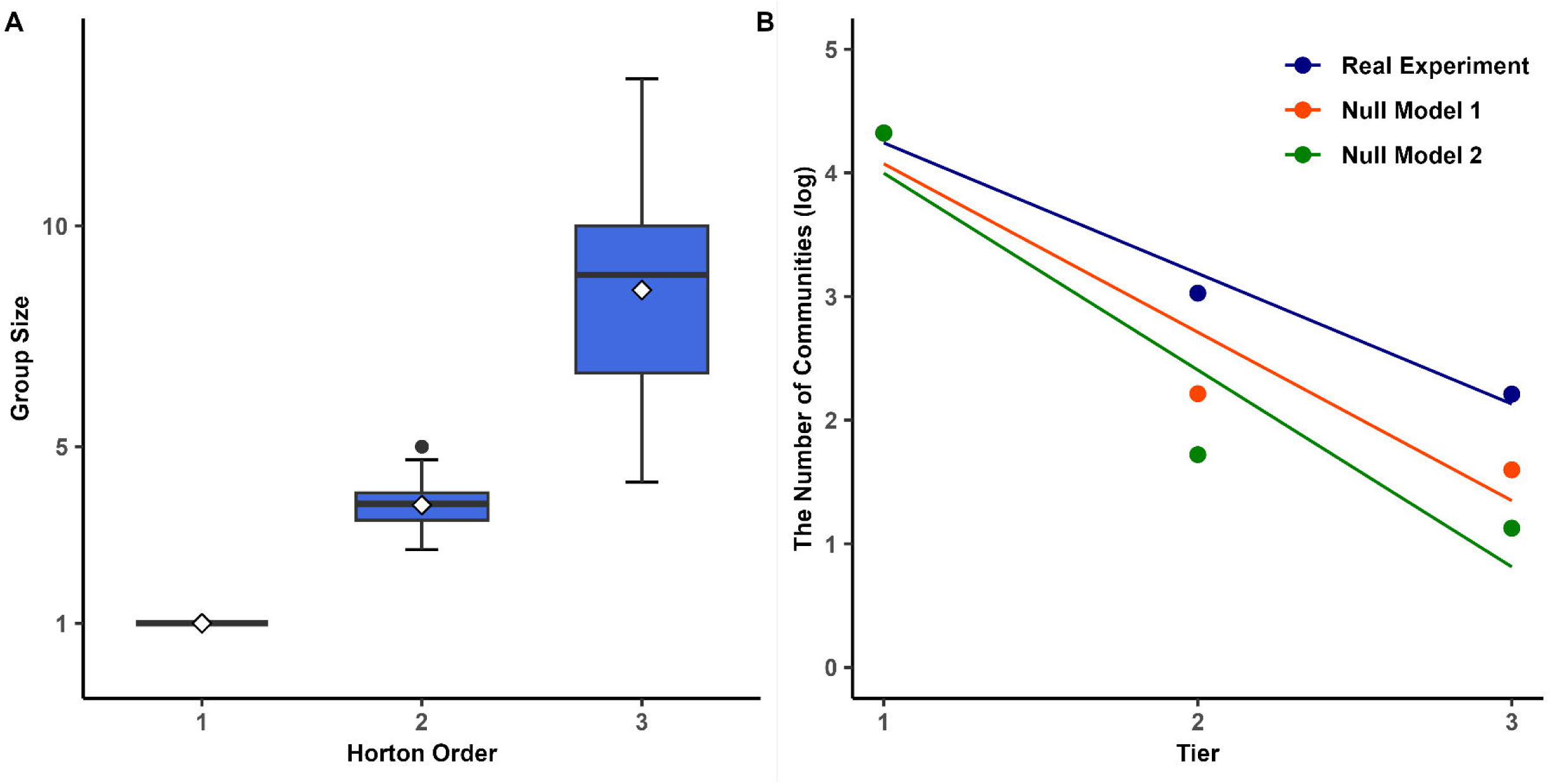
The community structure of zebra finch colony-level social networks contains multiple distinct tiers. (A) The distribution (box plots) and mean (diamonds) group sizes for different Horton orders (i.e. community tier in the social network). (B) The log number of groups in different Horton orders found in each day’s network (N = 30 days across four replicate colonies, bars show maximum and minimum values) among the real experient and two null models. The points are the logarithmic mean value of the number of relationships (note that all three datapoints overlap at x=1).

**Table 3.**
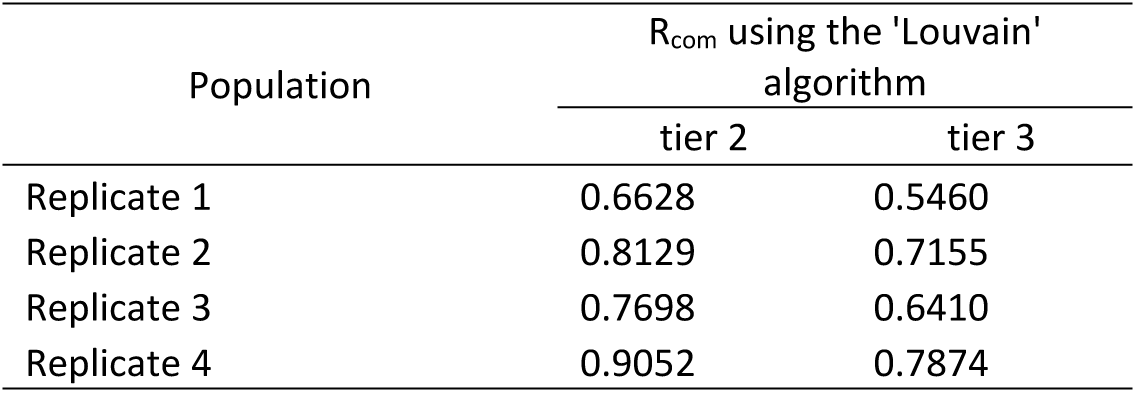
Revealing the multitiered social structure of zebra finch colonies. The robustness index (Rcom) suggests that a network has a substantial community structure when it exceeds 0.5. All colonies showed substantial structure for both tiers (range 0.55–0.91, note tier 1 is always the individual itself).

Two tiers were identified in both null models 1 and 2 (Fig. 2B), but the community structure in each colony was only robust in null model 1 (table S4). Like with the egocentric networks, we found that the tier corresponding to closest associates was absent in both null models, resulting in a steeper decline in the number of communities detected as a function of the number of tiers (Fig. 2B). In other words, there were more individuals within each group for a given tier in the null models than in the real, observed data. These results suggest that, as with the egocentric network approach, tiers representing weaker relationships (more occasional social partners) of the network can emerge as an outcome of spatial preferences, but the tiers representing stronger social relationships in the network emerge as a function of individuals’ specific social choices within the set of frequently available associates.

Finally, we used the Horton analysis (*54*) to test whether the community structure was also fractal-like. The regression slope between the community tier (the Horton number) and the log number of groups detected (calculated for each daily network) was -1.056 (Fig. 2B), corresponding to a Horton-Strahler branching ratio of 2.87 (see Methods for details). The linear fit of this regression suggests that a network with B branches can be split into B further branches, each growing in size by a factor of 2.87. These numbers align with the results of our analysis of the egocentric networks (Fig. 1E).

## Discussion

We found that the distribution of social contacts among zebra fiches is highly structured, both from the perspective of an individual and of the overall structure of the social network. Across four replicated colonies, individuals had, on average, 1-2 closest, 6-7 close, and 22-24 strong social partners, where social relationships were defined as sitting in close body contact. These patterns are qualitatively similar to those observed in humans (see Table 2 in Dunbar 2020 (*16*)), and the Horton branching ratio of 2.87 is similar to humans, primates, and other large mammals that all have values close to 3 (*54*). Importantly, the two null models showed that the properties that we detected were not attributed solely to differences in space use among individuals and reflected choosiness among individuals, even among their frequent social partners. Thus, zebra finches appear to maintain consistent higher-tier structure (strong social associates) across days that emerge from their choice of where to be while expressing notable carry-over of their closest social partners from day to day that emerged from social choices.

One important question is whether the tiered nature of the zebra finch social structure corresponds to a multilevel society. While individual zebra finches maintain a consistent set of closest associates, and these are also closest associates with one another (hence the fractal nature of the network), there is no evidence that these sets of close associates act as a stable social units. For example, the cosine similarities, though higher than expected by chance, never reached 1 (table S2). Multilevel societies have a distinct form of social structure where the higher tiers (e.g. close and strong social partners) are the result of interactions between groups of individuals that form lower tiers (the closest social partners), and thus where the lower tiers (or lowest tier) represent a distinct and stable social unit (*31*). For example, in superb fairywrens (*Malurus cyaneus*), members of the same social unit (called a breeding group) are always found together (*27*). Similarly, outside of breeding, groups of vulturine guineafowl remain together and move cohesively (*58*), even when there is a conflict over where or when to move (*59*). Thus, higher-tier associations in these species are the outcome of lower-tier collective decisions. By contrast, individual zebra finches are likely to have substantially greater independence when making decisions, meaning that they can temporarily fission and fusion as individuals, even if they do maintain several closest social relationships.

The pattern of multilevel social structure has been observed in many species, comprising levels such as troops, clans, and bands (Table 2). Previous studies have suggested that certain ecological pressures may contribute to the formation of these multilevel societies. These pressures include adapting to fluctuating food availability, maintaining spatial cohesion to avoid predator pressure (*60*), and fostering cooperative behaviour (*27*). While zebra finches do not live in multilevel societies, we found that the number of contacts increases exponentially in each tier of individuals’ social environments in a similar way.

These patterns raise questions about how multilevel societies evolve. While most research has focused on factors that drive between-group social contact, it is also possible that the multitied structure of multilevel societies could arise first. The resulting patterns, which include repeated social contacts among small subsets of individuals (closest relationships), may then promote various forms of cooperative investment [e.g. generalized reciprocity (*61*) or high-cost investments (*62*)]. The resulting interdependence among individuals (*63*) may then select for increased social stability, eventually leading to the formation of stable groups.

A major gap in knowledge is the question of how common tiered egocentric networks are in nature. Our literature search (Table 1) found relatively few examples, which were mostly in mammals. One reason for this gap is that it is progressively more challenging to collect the necessary data to accurately characterise each edge in a social network as we zoom out to higher tiers [see (*64*) for further discussion]. Without the ability to accurately estimate each edge, subsequent analyses aiming to identify putative tiers would become subject to substantial error. In our study, we were able to identify four putative tiers in zebra finch social networks because our tracking system allowed us to collect high-resolution spatial and temporal data in much larger colonies than what had been previously studied. Previous to our study, individuals had been studied in social contexts where they were unlikely to face many of the limitations that give rise to the patterns we uncovered (e.g. in small colonies) or could only track a small portion of the population (*43*). Yet, even our colonies, which were comprised of 80 birds each, were also much smaller than some of the flock sizes naturally found in nature. For example, zebra finches can exist in flocks of up to 300 birds in non- breeding seasons (*65*). Thus, we were likely to have been limited in our ability to quantify the precise number of individuals present within the higher social tiers of individuals’ egocentric networks (i.e. the fourth tier or any tiers above that). Studying fine-scale structures in such large groups remains a long-term challenge for the field (*33*). Future work on larger colonies would be needed to better resolve the higher tiers of zebra finch societies.

A major question is where the outer bound to individuals’ abilities to maintain known social relationships is. Some bird species, such as superb starlings (*Lamprotornis superbus*) and vulturine guineafowls (*Acryllium vulturinum*), can maintain large group sizes (e.g. up to at least 65 individuals in the latter) (*66*, *67*) with consistent social membership that can last over months or years (*68*). Thus, it is likely that—at least in some birds—the outer bound could exceed 60 individuals. From the regression in Fig. 2B, we can also find that the horizontal intercept in the observed data is about 5, meaning that zebra finches may have the potential to form a community of the fifth Horton order. The size of a community at that order would be about 68 individuals, which is too large for us to have been able to detect statistically (i.e. it would require studying colonies of > 120 birds). Previous studies (*41*) also found that the mean size of basic social units in wild zebra finches is 2.9, which is similar to the size of our inferred second tier communities (i.e. closest relationships). The match between the pattern observed in the wild and those in our experiment may also reflect the fact that groups in zebra finches are primarily organised around mated pairs, which maintain close spatial and temporal associations, and that these often associate with other non- breeding social partners (*43*). Together, these insights suggest that the colonies in our studies can effectively reflect the actual social structure in zebra finches, at least in the communities capturing lower tiers.

Our estimates of the number of closest social contacts for zebra finches are consistent with those in other species (e.g. Table 1 and in humans), but the number of close and strong social contacts is marginally higher. This could be due to several methodological differences. One potential reason is that we kept individuals in high densities, and higher densities typically lead to higher rates of social encounters (*69*). However, as we normalised each individual’s distribution of edge weights (see Methods), we expected that density should not have a large effect because we could characterise properties that correspond to the shape of the distribution rather than the absolute contact rates themselves. Future studies could investigate the sensitivity of the estimates to population density and the robustness of drawing absolute estimates from studies with very different means of collecting the underlying social data.

We demonstrated that zebra finches’ social networks contain distinct tiers and found a scaling ratio by Horton analysis of 2.87. Previous studies had found similar scaling ratios in humans [3.11 (*16*)] and social animals [2.99 for elephants, 3.05 for geladas, 3.04 for hamadryas and 3.80 for orcas (*54*)] and suggested that scaling ratios of around 3 may be a universal feature of multilevel animal societies (*54*). A key question is whether social animals evolve to have some convergent structural patterns in their social contacts, such as the differentiation of the egocentric relationships and the stratification of the society or some other forms of fractal patterns, because such structures are adaptive (*sensu* (*70*)) or whether this scaling ratio captures a fundamental property of how animal societies self-organise. For example, such patterns could emerge because social animals face similar constraints of time and cognition (*71*, *72*). Previous work in zebra finches found that rates of clumping increased with temperature (*47*), suggesting that such behaviour is time-limited. There is also increasing awareness of the role that space use plays in shaping the overall structure of social networks (*39*), with previous work on a population-level network of great tits (*Parus major*) highlighting the clustering of individuals in space as generating a pattern that can appear like a multilevel network. In our study, we could use null models to explicitly disentangle the contribution of spatial and social processes, confirming the importance of space in determining who is available to flock with (higher tiers) and the importance of social choices in refining associations to a smaller subset of individuals (lower tiers) within the local social environment.

## Materials & Methods

### Experiment model and data collection

Between December 2017 and March 2018, we conducted continuous and fine-scale tracking in four large replicated colonies (N≍80, see table S1; the dimension of each aviary: 5m x 2.0m x 2.5m) of zebra finches. The zebra finches were placed in mixed-sex colonies (e.g. N=40 males and 40 females) for 30 days. In replicates 1 and 2 (commencing December 2017), the birds were 170 ± 25 days old at the start of the experiment, with an age range of 105- 199 days; in replicates 3 and 4 (commencing January 2018), the birds were 200 ± 29 days old, with an age range of 120-241 days. During the experiment, we used an automated barcode-based tracking system to record the positions of individuals (*46*). Here we report the result from the mixed-sex colonies.

Each individual was fitted with a unique machine-readable barcode (*51*). In each aviary, eight cameras (8-megapixel Camera Module V2; RS Components Ltd and Allied Electronics Inc.), each connected to a Raspberry Pi (Raspberry Pi 3 Model Bs; Raspberry Pi Foundation), were strategically placed. These cameras recorded individuals at six perches and at two feeders.

Between 05:30 and 20:00, when lights were switched on, each camera took a picture every two seconds. The structure and size of the perches and feeders in each aviary were designed to allow individuals to freely interact with others or move between groups at their discretion.

We first defined social connectivity among individuals based on spatial proximity, defining two individuals to be closely associated and likely interacting if they are in physical contact [’clumping’ (*37*)]. Zebra finches often maintain this body contact for extended time periods, during which they often engage in allopreening and other affiliative interactions (*45*). Body contact was defined using a threshold distance of 80 pixels between two barcodes (fig. S1), which corresponds to the width of a zebra finch at the height at which the cameras were mounted (*37*). Using the second-by-second proximity data, we then calculated the propensity for each pair of zebra finch to be in physical contact on each day, using a modified simple ratio index [the total number of detections together divided by the number of frames in which both were observed on the same perch (*73*)]. This edge definition captures individuals’ social decisions in terms of whom to have contact with independently of spatial drivers that determine their opportunity to interact [i.e., spatial proximity (*74*)].

### Detection of relationship tiers

We used the Jenks natural breaks algorithm (*55*), which is implemented in R package BAMMtools (*75*), to identify different categories of social contacts in the egocentric networks of 152 male and 151 female zebra finches. This algorithm identifies clusters (sets of relationships with different edge weights) by minimising within-cluster distances and maximising between cluster distances. We calculated the goodness of variance fit index (GVF) for the Jenks natural breaks algorithm with different numbers of clusters:

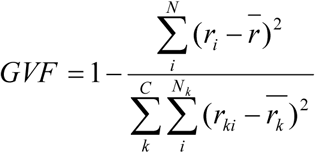

where *C* is the number of clusters, *N* is the number of all individuals in the ego-networks, *N_k_* is the size of the kth cluster identified by the Jenks natrual breaks algorithm, *r_ki_* is the normalised contact rate with the focal bird of the ith individual in the kth cluster, where the normalised contact rate means the contact rate of i to the focal bird divided by the mean contact rate of the focal bird with all other colony members, and 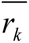 means the mean nomalised contact rate in the kth cluster. The GVF varies between 0 and 1, and gives a measure of minimising the variance within classes and maximising the variance between classes when the data are partitioned into N clusters. As the GVF increases with the number of clusters, we should use the cut-off at GVF = 0.85 recommended by Coulson (*56*) to identify the optimal number of clusters. We then found the optimal clustering number in all networks, which is four (Fig. 1C), to get the differentiated relationships. For an individual’s egocentric networks, the number of closest relationships is defined as the size of the first cluster; the number of close relationships is the sum of the sizes of the first and second clusters, corresponding to human friendships; the number of strong relationships is the sum of the first three clusters; and the last cluster is classified as acquaintances, which is not seen as a special social relationship. Then, we first made a comparison between the mean number of relationships among males and females using mixed-effect linear regression by *lme4* package in r (*76*). In this analysis, sex was treated as a fixed effect while individual identity was considered as a random effect. Subsequently, we tested the difference between same-sex and opposite-sex relationships within and among sexes in the same way. To quantify the stability of the identities of the social partners across days, we used the cosine similarity, which is defined as:

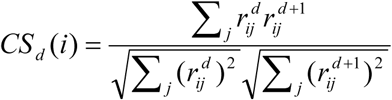

where 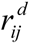 represents the normalise contact rate of i and j on day d, which is the contact rate of i and j on day d divided by the mean contact rate of i with all other colony members on that day. The cosine similarity ranges from 0 to 1, with *CS_d_* (*i*) being high if the contact rate with all social partners was similar across days (e.g. the close relationships of i on day d are still close relationships on day d+1). *CS_d_* (*i*) is low if the contact rate change from day to day. To get a better understanding of the value obtained, we performed 1000 permutations of the identities of individuals in each daily networks (independently) to generate a null distribution of the cosine similarity expected by chance.

To further illustrate the fractal structure present in the egocentric networks, which means the number of relationships increases exponentially across layers, we used the Horton analysis (further details provided in the subsequent section). A linear mixed-effect regression model was constructed by *lme4* (*76*), with the log number of relationships as dependent variables, tier as a fixed effects, and individual identity as a random effect. The existence of the fractal structure can be validated by the presence of a notable linear relation.

### Null models

We generated the null model 1 by permuting the individual identities on the same perch for every record within the original second-by-second proximity data, and the null model 2 by pair of zebra finch to be in physical contact on each day, using a modified simple ratio index similar to our approach in the real situations. The relationships of different types were recognised by using the thresholds determined by Jenks natural breaks algorithm in the real experiments, which are more likely to accurately represent breaks in different relationships in the real situation. This method allowed for a clearer comparison between the real experiments and the null models. The techniques used to determine the numbers of relationships and the pattern of increase across tiers are same as those employed in the real experiments.

### Extracting community structures

To determine whether zebra finch social networks have a tiered structure, we started by constructing one social network for the duration of each colony. To get the different tiers of community structure for each colony, we used the ’Louvain’ algorithm from *igraph* (*77*). This algorithm iteratively identifies the hierarchical subgroups and maximises the connection density within these subgroups compared to between them at different tiers (*31*), and it revealed second and third tier communities (the first tier being each single individual). We then used the resampling algorithm described by Shizuka and Farine (*78*) to estimate the robustness of these communities in both detected tiers by generating a network of community co-membership.

While there has been some debate about the use of bootstrapping methods as a means to control for dependencies in social network data (*79*), it remains a valuable approach to test the robustness of the communities. In brief, we created 1000 networks using bootstrapped versions of our observation dataset (resampling the second-by-second data with replacement, keeping all individuals in the network). In each of these networks, we used the same ’Louvain’ algorithm to determine the membership of individuals into communities. Our community co-membership networks reflect the proportion of these networks in which each pair (dyad) of individuals co-occurred in the same community. Finally, we extracted the community structure of this network, and measured the assortativity [using *assortnet* (*80*)] of the network according to the community structure. The resulting value (R_com_) reflects the partitioning of the network into substantial community structure when values exceed 0.5, and a lack of community structure when below 0.5.

We then used Horton analysis to calculate the Horton-Strahler branching ratio (*81*) (B) between the three tiers in the real observed data and two null models:

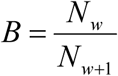

where the *N_w_* is the number of groups in order w detected using the ’Louvain’ algorithm in each daily networks, the equation can be rearranged on the log scale as:

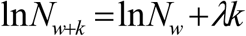

where *λ*= −ln *B* and *k* = Δ*w*. We fitted the equation to the log number of the frequency of the communities found in each daily networks by linear mixed-effect regression using *lme4*, and the rooms and replicates are included as random effects, to get the regression slope *λ*, which can be then used to calculate the Horton-Strahler branching ratio by *B* = *e*^−^*^λ^*.

## Acknowledgments

We thank Wolfgang Forstmeier and Bart Kemperneners for supporting study species and facilities.

## Author Contributions

Conceptualization: DRF, DW

Methodology: YZ, DRF, DW

Investigation: YZ, DRF, DW

Analysis: YZ, DRF, DW

Supervision: DRF, DW

Writing—original draft: DRF

Writing—review & editing: YZ, XJ, LMA, DW, DRF

## Data and Materials Availability

All data needed to evaluate the conclusions in the paper are present in the paper and/or the Supplementary Materials. (http://datadryad.org/stash/share/rT8w0Ngkn3Lw6GARfJxLfqn9UqrwkiPCZYVXzY7xooM)

## Supplement Materials

This PDF file includes: Figs. S1 to S6 Tables S1 to S4 Movie S1

