## Supplement Material for "Social and spatial drivers of the multitiered structure of zebra finch social networks"

This PDF file includes:

Figs. S1 to S6

Tables S1 to S4

Movie S1

**Table S1. The total number of birds and the number of each sex in different populations.**

| Population | N | $N_{male}$ | $N_{female}$ |
|---|---|---|---|
| Replicate 1 | 80 | 40 | 40 |
| Replicate 2 | 80 | 40 | 40 |
| Replicate 3 | 80 | 40 | 40 |
| Replicate 4 | 63 | 32 | 31 |

**Table S2. The average cosine similarity (N = 303) across 30 days.** This range is substantially higher than what is expected by chance (0.19-0.35), suggesting that even shortly after being introduced to unfamiliar birds, zebra finches develop and maintain consistent preferred social bonds.

| Day | Cosine Similarity |
|---|---|
| 1 | 0.6958 |
| 2 | 0.6972 |
| 3 | 0.7018 |
| 4 | 0.7332 |
| 5 | 0.7315 |
| 6 | 0.7253 |
| 7 | 0.7280 |
| 8 | 0.7407 |
| 9 | 0.7313 |
| 10 | 0.7390 |
| 11 | 0.7550 |
| 12 | 0.7478 |
| 13 | 0.7422 |
| 14 | 0.7202 |
| 15 | 0.7106 |
| 16 | 0.7216 |
| 17 | 0.7276 |
| 18 | 0.7246 |
| 19 | 0.7321 |
| 20 | 0.7130 |
| 21 | 0.6754 |
| 22 | 0.7042 |
| 23 | 0.7036 |
| 24 | 0.6952 |
| 25 | 0.7192 |
| 26 | 0.7011 |
| 27 | 0.7149 |
| 28 | 0.7239 |
| 29 | 0.6689 |

**Table S3. The average group size of communities detected by the 'Louvain' algorithm in every day's network.**

| Day | Level 2 | | | | Level 3 | | | |
|---|---|---|---|---|---|---|---|---|
| | Replicate 1 | Replicate 2 | Replicate 3 | Replicate 4 | Replicate 1 | Replicate 2 | Replicate 3 | Replicate 4 |
| 1 | 4.71 | 3.81 | 4.21 | 3.50 | 13.33 | 8.89 | 11.43 | 9.00 |
| 2 | 3.81 | 3.81 | 4.21 | 3.94 | 11.43 | 8.89 | 10.00 | 9.00 |
| 3 | 4.00 | 3.81 | 4.00 | 3.71 | 11.43 | 8.00 | 10.00 | 7.00 |
| 4 | 3.81 | 3.33 | 3.64 | 3.94 | 11.43 | 7.27 | 10.00 | 7.00 |
| 5 | 5.00 | 3.64 | 4.71 | 3.71 | 11.43 | 8.89 | 10.00 | 7.88 |
| 6 | 3.64 | 3.20 | 4.00 | 3.71 | 8.89 | 6.67 | 10.00 | 9.00 |
| 7 | 5.00 | 3.64 | 4.44 | 3.94 | 10.00 | 8.00 | 11.43 | 7.00 |
| 8 | 4.21 | 3.81 | 3.48 | 3.94 | 10.00 | 7.27 | 10.00 | 7.00 |
| 9 | 3.81 | 3.81 | 4.00 | 3.94 | 10.00 | 7.27 | 11.43 | 5.73 |
| 10 | 4.71 | 4.00 | 3.20 | 3.71 | 13.33 | 8.00 | 11.43 | 5.73 |
| 11 | 4.21 | 3.48 | 4.00 | 3.50 | 11.43 | 8.89 | 10.00 | 5.73 |
| 12 | 4.21 | 3.81 | 4.21 | 3.50 | 13.33 | 6.67 | 8.89 | 5.73 |
| 13 | 4.00 | 3.48 | 3.81 | 3.15 | 13.33 | 6.67 | 11.43 | 6.30 |
| 14 | 4.00 | 3.48 | 3.81 | 3.50 | 8.89 | 8.00 | 11.43 | 6.30 |
| 15 | 4.21 | 3.33 | 4.00 | 3.32 | 10.00 | 8.00 | 10.00 | 6.30 |
| 16 | 3.48 | 3.33 | 3.81 | 3.32 | 10.00 | 8.00 | 11.43 | 6.30 |
| 17 | 3.64 | 3.48 | 3.20 | 3.15 | 7.27 | 6.67 | 8.89 | 6.30 |
| 18 | 3.81 | 3.20 | 4.00 | 2.86 | 11.43 | 7.27 | 11.43 | 5.25 |
| 19 | 3.81 | 2.96 | 3.64 | 3.32 | 11.43 | 7.27 | 8.89 | 6.30 |
| 20 | 4.44 | 3.33 | 3.48 | 3.32 | 8.89 | 8.00 | 10.00 | 5.73 |
| 21 | 4.44 | 3.81 | 3.33 | 3.00 | 11.43 | 7.27 | 6.67 | 5.73 |
| 22 | 4.00 | 3.33 | 3.08 | 3.00 | 11.43 | 6.67 | 8.89 | 5.25 |
| 23 | 3.48 | 3.64 | 3.08 | 3.32 | 10.00 | 8.89 | 8.00 | 5.25 |
| 24 | 4.00 | 3.33 | 3.81 | 3.15 | 8.89 | 8.00 | 10.00 | 4.50 |
| 25 | 4.00 | 3.33 | 3.48 | 3.00 | 8.89 | 5.71 | 10.00 | 4.20 |
| 26 | 3.81 | 3.64 | 3.81 | 3.00 | 8.89 | 6.67 | 8.89 | 5.25 |
| 27 | 3.81 | 3.81 | 3.64 | 3.15 | 8.89 | 7.27 | 10.00 | 4.85 |
| 28 | 4.21 | 2.96 | 3.48 | 2.86 | 11.43 | 6.67 | 8.00 | 5.73 |
| 29 | 3.81 | 2.67 | 3.81 | 3.00 | 8.00 | 6.67 | 11.43 | 5.25 |
| 30 | 4.44 | 3.20 | 3.64 | 3.00 | 10.00 | 6.15 | 10.00 | 5.25 |

**Table S4. The robust index ($R_{com}$) in the two null models.** The index exceeds 0.5 in most cases of null model 1, which suggests a substantial community structure. However, in null model 2, the $R_{com}$ cannot be calculated. This is because, although community structures can be identified in each day's network, they cannot be detected when applied to the mean clumping rate network of the 30-day period and the networks generated by the bootstrap method (i.e. all individuals are allocated into one group). This indicates that the community is not stable in null model 2.

| Population | $R_{com}$ using the 'Louvain' algorithm | |
|---|---|---|
| | level 2 | level 3 |
| Null model 1 replicate 1 | 0.6123 | 0.6403 |
| Null model 1 replicate 2 | 0.6963 | 0.6241 |
| Null model 1 replicate 3 | 0.6902 | 0.5939 |
| Null model 1 replicate 4 | 0.4102 | 0.6154 |
| Null model 2 replicate 1 | NA | NA |
| Null model 2 replicate 2 | NA | NA |
| Null model 2 replicate 3 | NA | NA |
| Null model 2 replicate 4 | NA | NA |

**Figure S1. Example tracking image of zebra finches in aviaries.** The birds enclosed in the red circle are categorized as "clumping" individuals. The distance between these birds is less than 80 pixels in the raw image, indicating that they are engaged in social interactions.

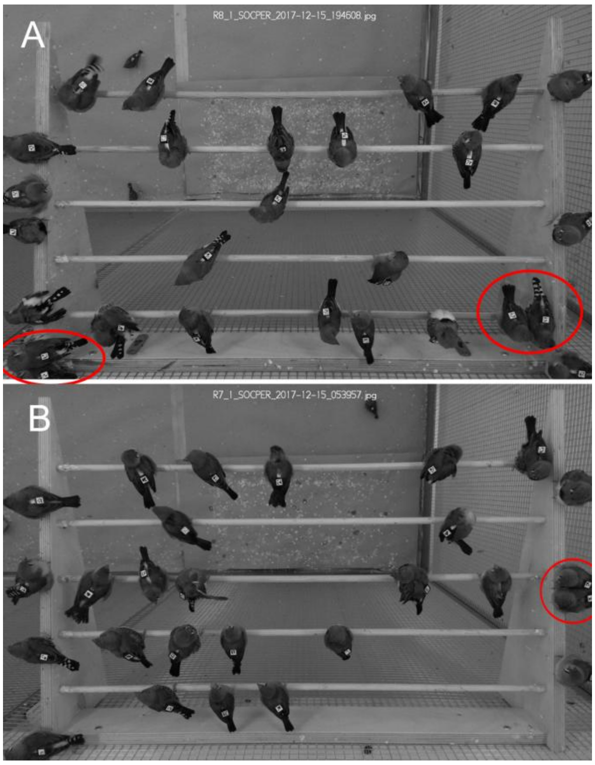

**Figure S2. The number of relationships in zebra finches** (A) The number of relationships detected using Jenks natural breaks algorithm for each sex (males: n = 152 and females: n = 151) in colonies. (B) Males and females showed differences in their propensity to have same- vs. opposite-sex relationships. Significant differences (two sample t-tests) between the number of relationships are shown as * p ≤ 0.05, ** p ≤ 0.001 and *** p ≤ 0.001.

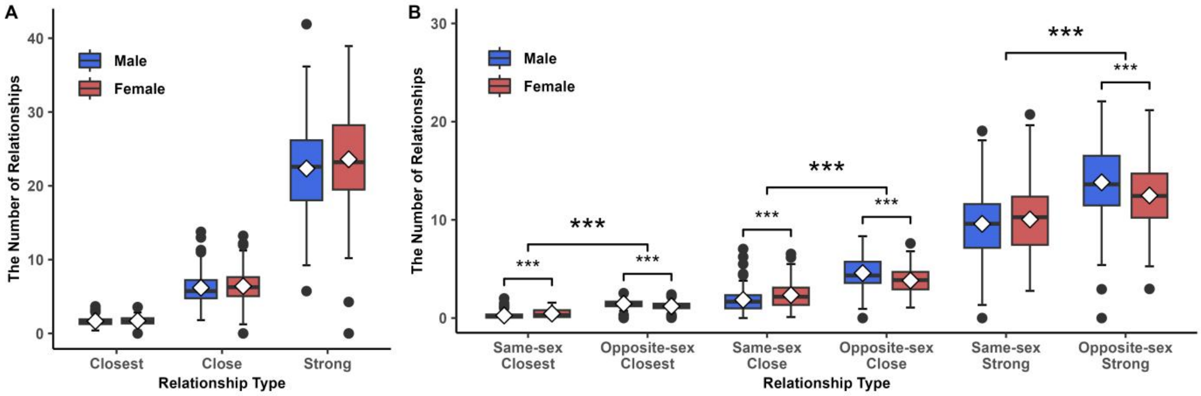

**Figure S3. All individuals' ego-network with the three layers of relationships in the replicate 1.**

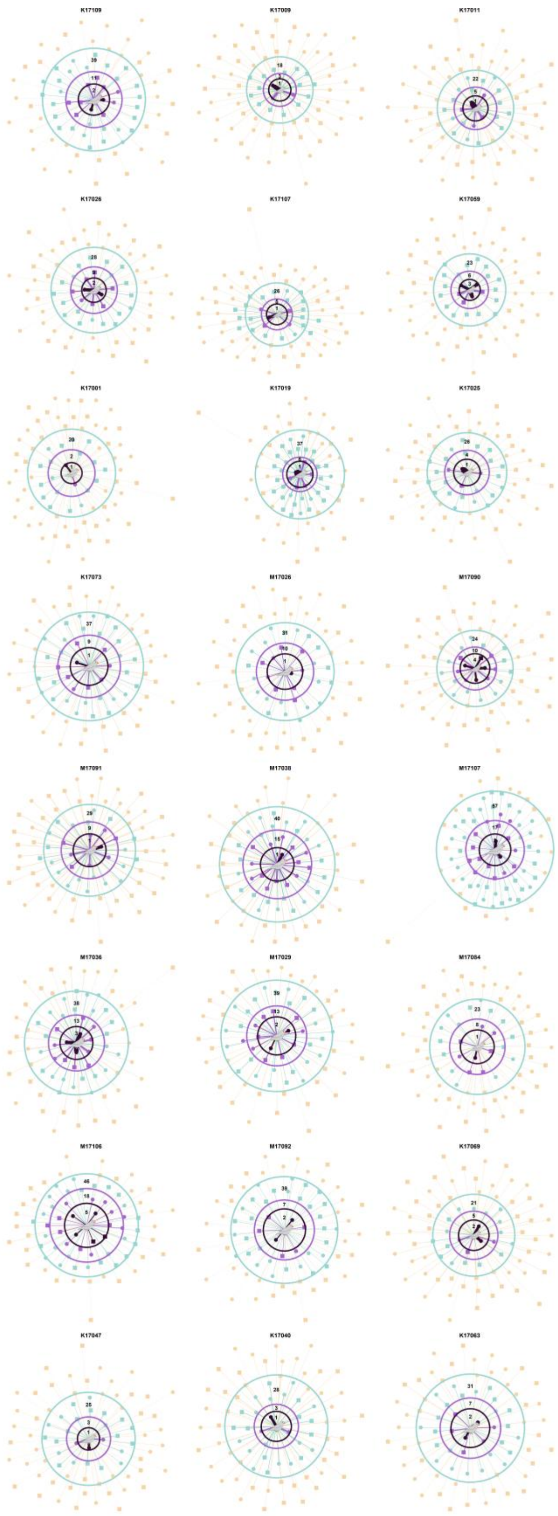

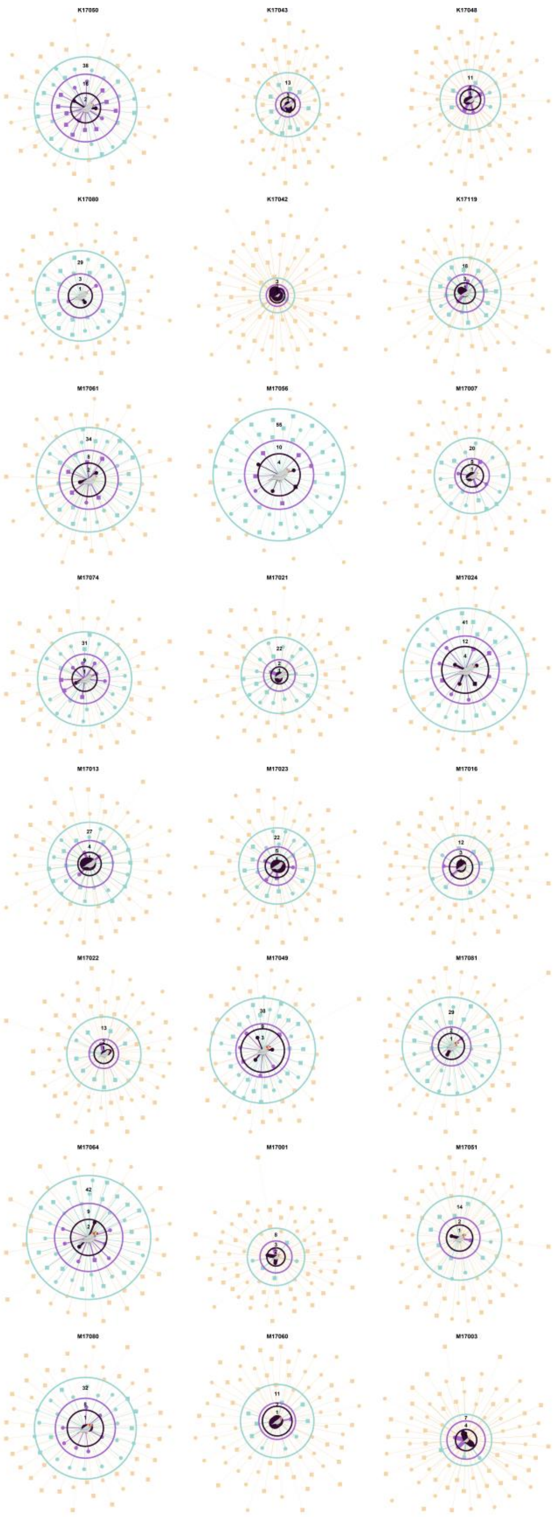
K17050
K17043
K17048
K17080
K17042
K17119
M17061
M17056
M17007
M17074
M17021
M17024
M17013
M17023
M17016
M17022
M17049
M17081
M17064
M17001
M17051
M17080
M17060
M17003

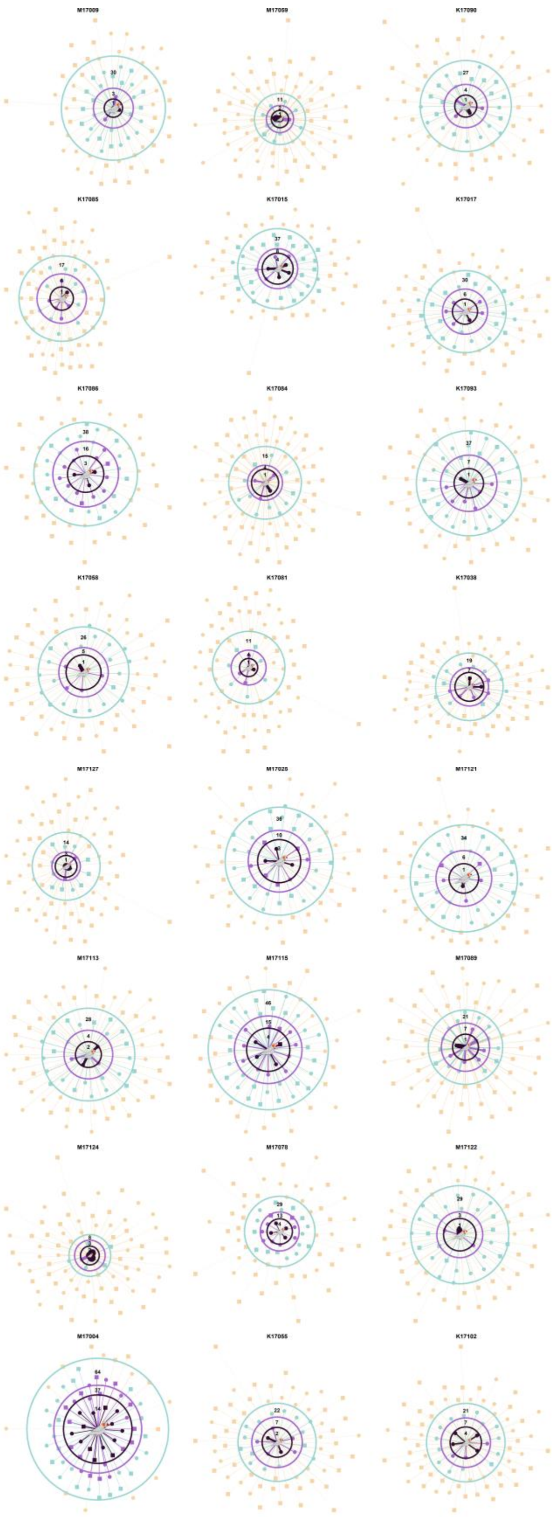
M17009
M17059
K17090
K17085
K17015
K17017
K17086
K17084
K17093
K17058
K17081
K17038
M17127
M17025
M17121
M17113
M17115
M17089
M17124
M17078
M17122
M17004
K17055
K17102

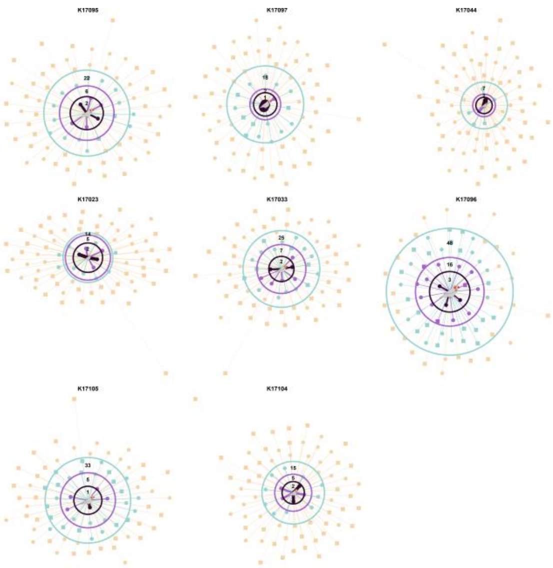
K17095
22
6
2
K17097
18
1
K17044
7
K17023
5
2
K17033
25
7
2
K17096
48
16
3
K17105
33
5
1
K17104
15
5
2

**Figure S4. All individuals' ego-network with the three layers of relationships in the replicate 2.**

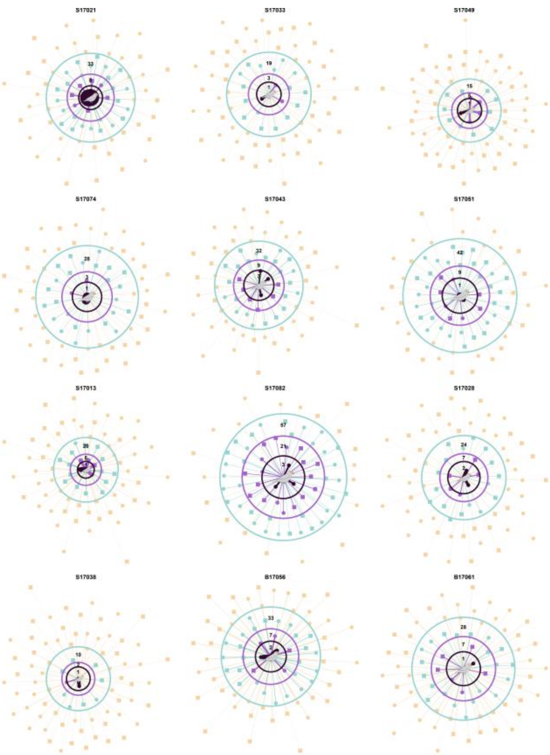


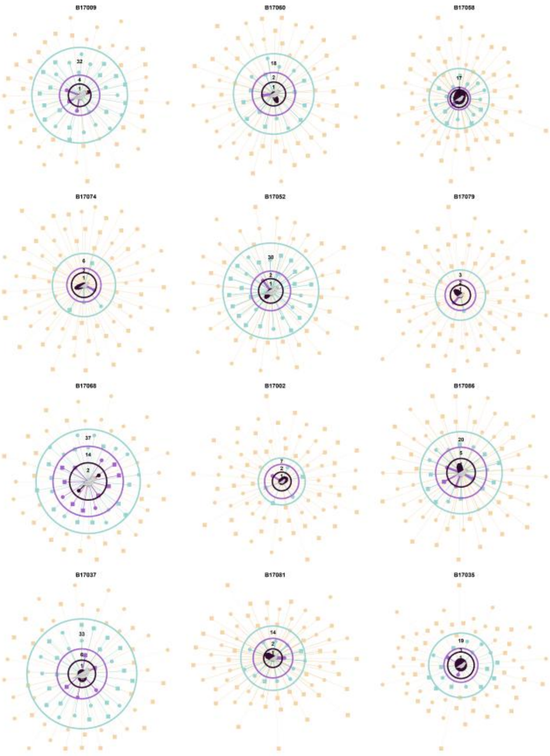

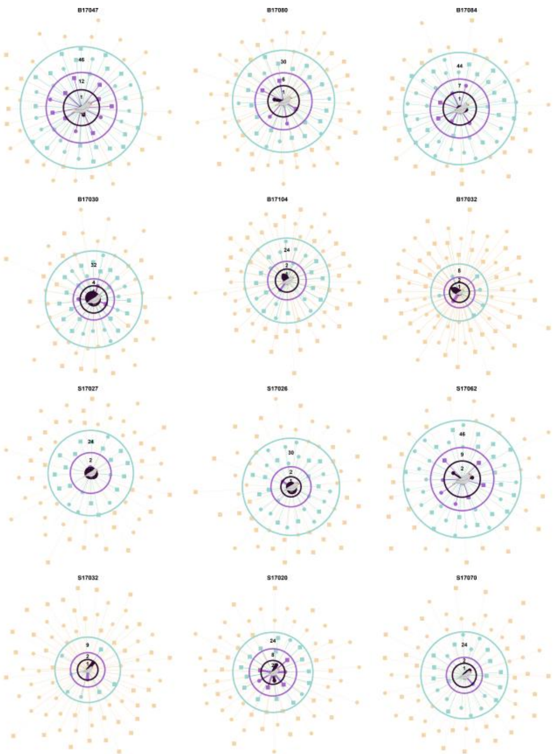
B17047
B17080
B17084
B17030
B17104
B17032
S17027
S17026
S17062
S17032
S17020
S17070

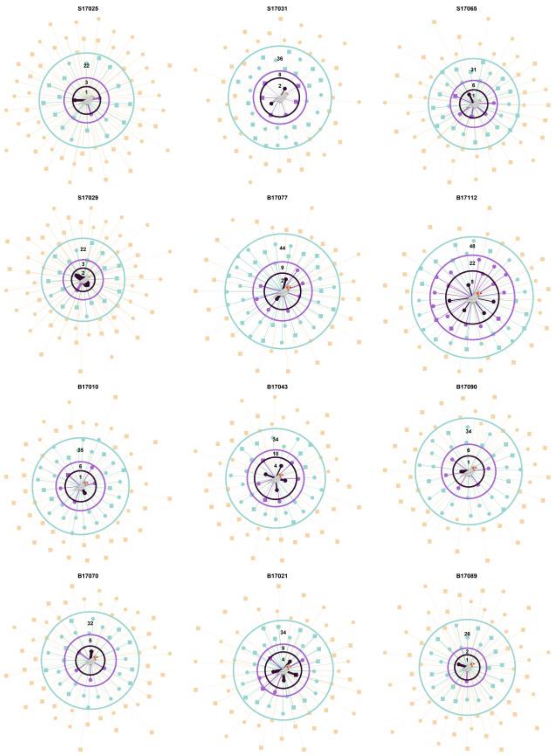
S17025
S17031
S17065
S17029
B17077
B17112
B17010
B17043
B17090
B17070
B17021
B17089

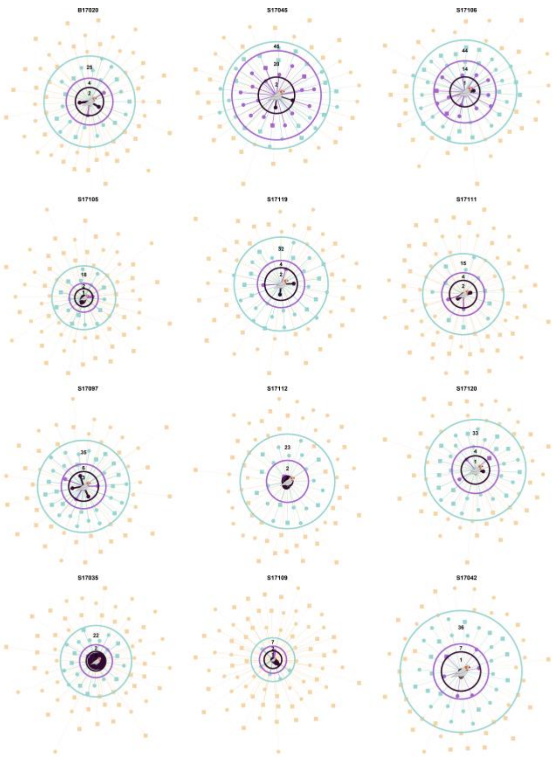
B17020
S17045
S17106
S17105
S17119
S17111
S17097
S17112
S17120
S17035
S17109
S17042

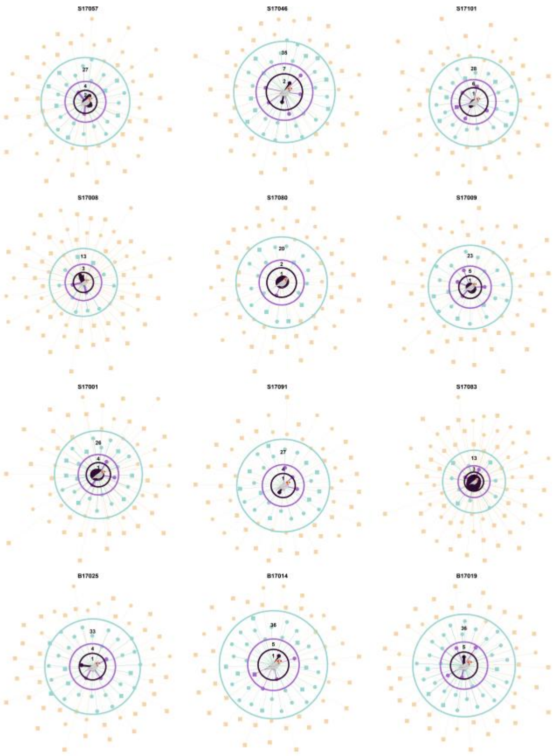
S17057
S17046
S17101
S17008
S17080
S17009
S17001
S17091
S17083
B17025
B17014
B17019

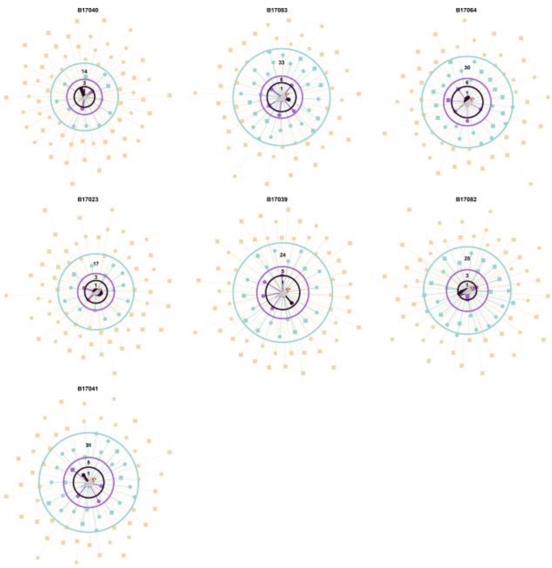
B17040
14
3
B17083
33
6
1
B17064
30
6
1
B17023
17
3
1
B17039
24
5
1
B17082
28
3
1
B17041
31
5
1

**Figure S5. All individuals' ego-network with the three layers of relationships in the replicate 3.**

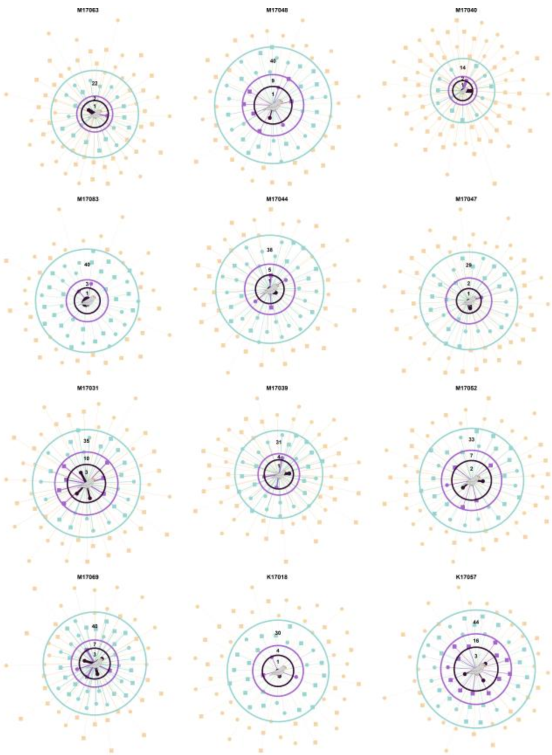

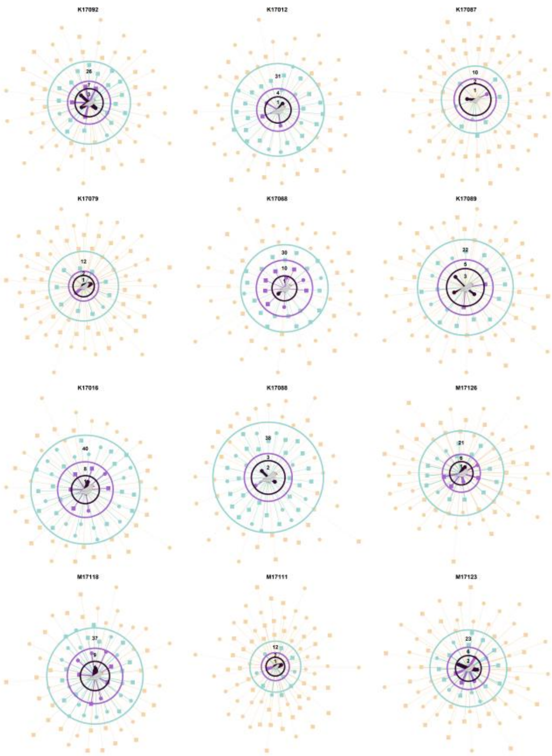

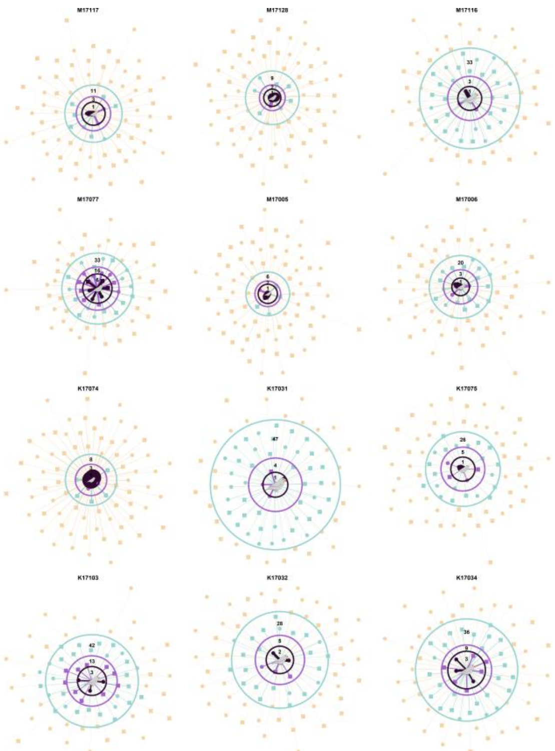
M17117
M17128
M17116
M17077
M17005
M17006
K17074
K17031
K17075
K17103
K17032
K17034

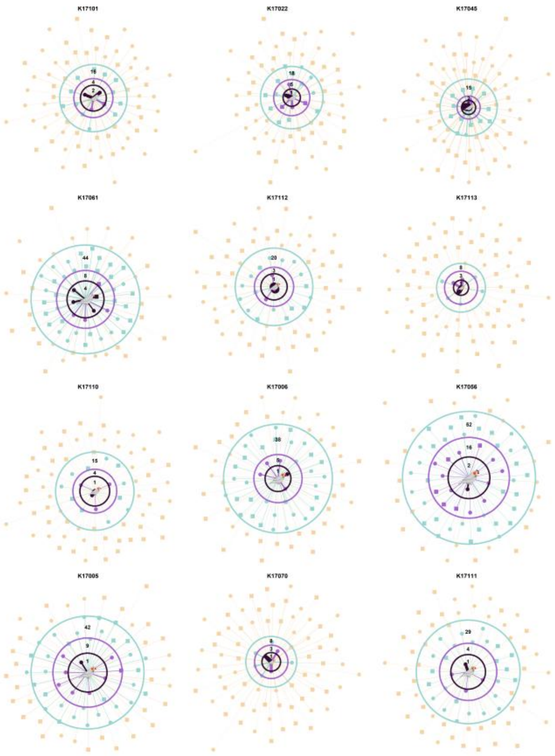
K17101
K17022
K17045
K17061
K17112
K17113
K17110
K17006
K17056
K17005
K17070
K17111

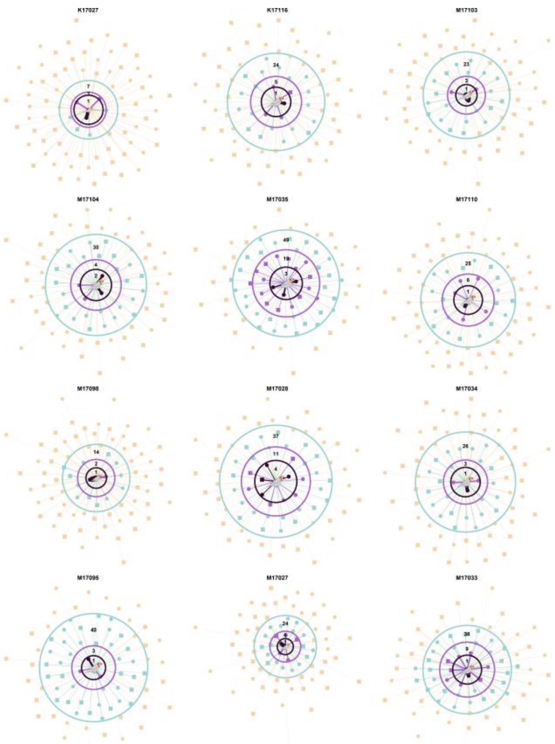

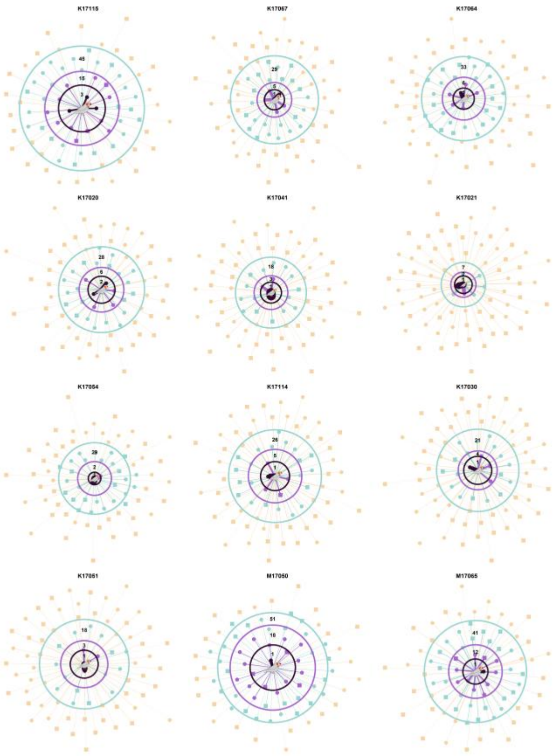

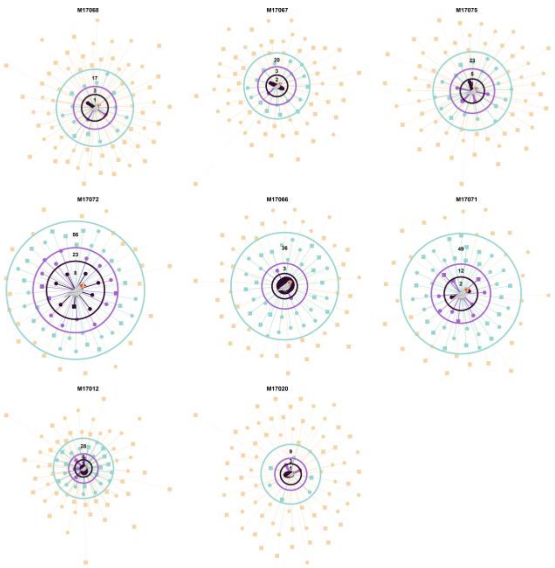
M17068
M17067
M17075
M17072
M17066
M17071
M17012
M17020

**Figure S6. All individuals' ego-network with the three layers of relationships in the replicate 4.**

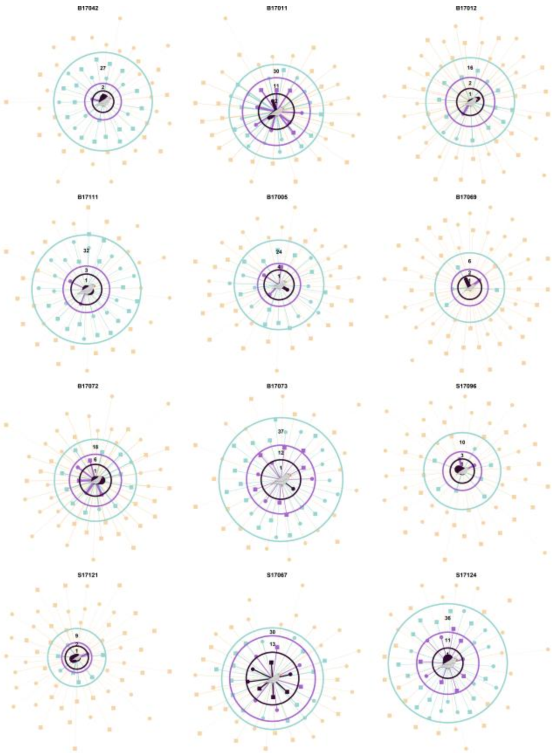

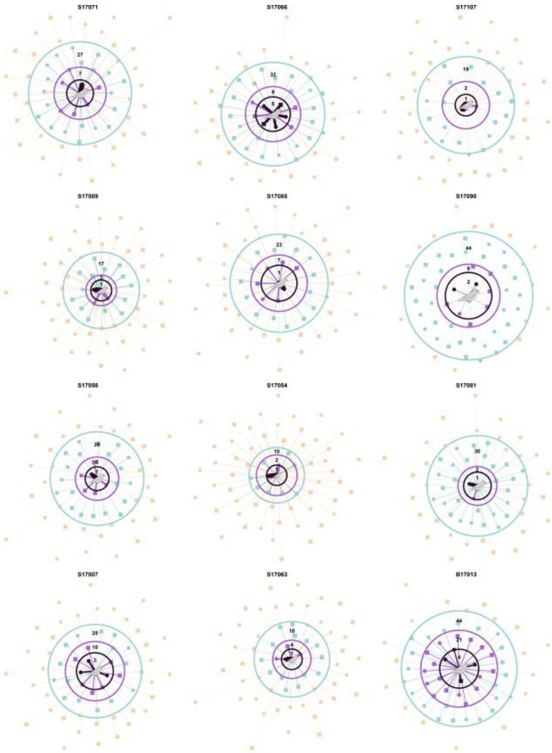

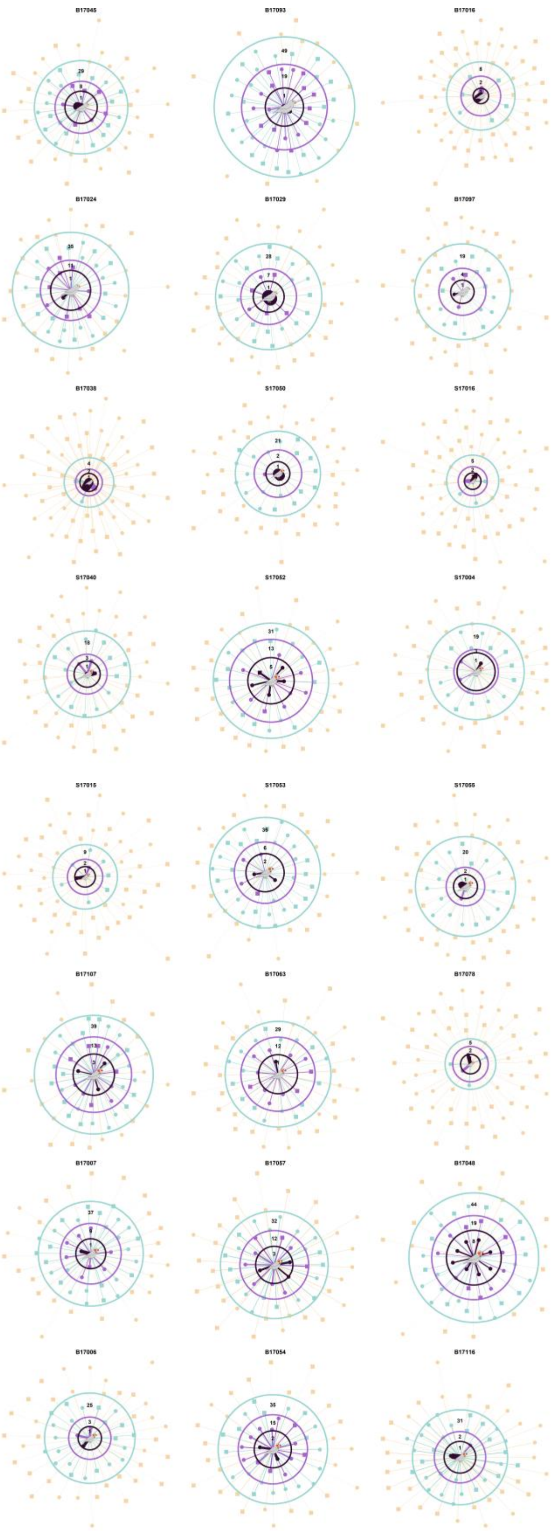
B17045
B17093
B17016
B17024
B17029
B17097
B17038
S17050
S17016
S17040
S17052
S17004
S17015
S17053
S17066
B17107
B17063
B17078
B17007
B17057
B17048
B17006
B17054
B17116

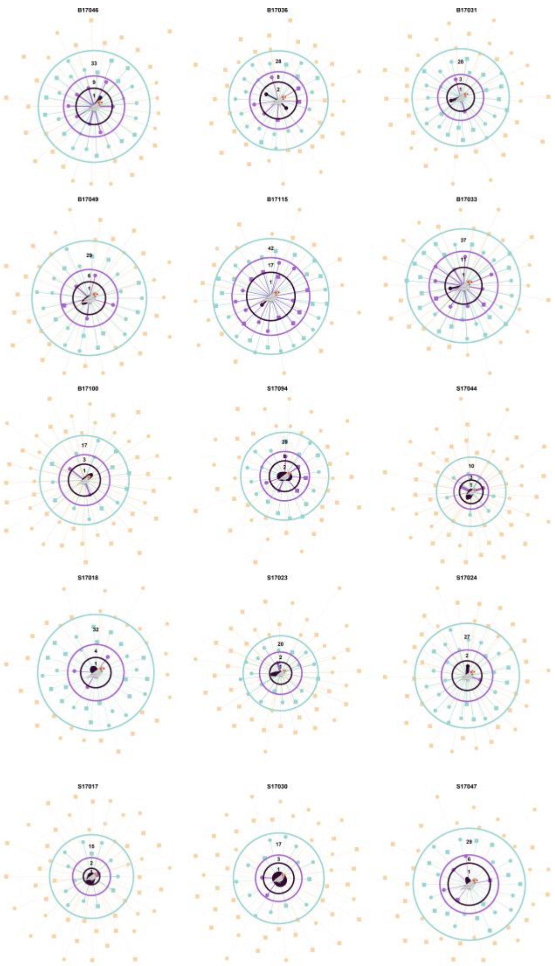
B17046
B17036
B17031
B17049
B17115
B17033
B17100
S17094
S17044
S17018
S17023
S17024
S17017
S17030
S17047

**Movie S1. Example tracking video of zebra finches in aviaries.** This example shows the progression of zebra finch activity on a perch.
